# SCOUT: Ornstein–Uhlenbeck modelling of gene expression evolution on single-cell lineage trees

**DOI:** 10.1101/2025.11.12.688020

**Authors:** Hannah Stuart, Aaron McKenna

## Abstract

Understanding the evolutionary dynamics of clonal populations is essential for uncovering the principles of development, disease progression, and therapeutic resistance. Recent advances in single-cell lineage tracing and transcriptomics enable such analyses by combining heritable barcodes with cell-state information. Here, we present SCOUT (<u>s</u>ingle-<u>c</u>ell <u>O</u>rnstein–<u>U</u>hlenbeck <u>t</u>rees), a framework that models gene expression dynamics along single-cell lineage trees using Ornstein–Uhlenbeck processes to distinguish neutral drift from selective pressure. Using simulations, we demonstrate that SCOUT accurately classifies genes based on their underlying evolutionary models. We further validate SCOUT in Caenorhabditis elegans development, identifying biological processes under selection across distinct developmental contexts. Finally, we apply SCOUT to a lung adenocarcinoma xenograft model, revealing key regulators of metastatic progression and tumor microenvironmental adaptation. By integrating lineage and transcriptomic data, SCOUT provides a powerful evolutionary lens for dissecting the forces that shape cell fate.

## Introduction

A number of recent publications have generated high-resolution, single-cell lineage trees of processes in both normal development and in diseases such as cancer (Quinn et al. 2021; Kalhor et al. 2018; Saxe et al. 2025; Simeonov et al. 2021; Li et al. 2025). Using CRISPR-based single-cell lineage tracing (scLT), these papers map the ancestral relationship of thousands of cells through time, jointly captured with their transcriptional state (McKenna and Gagnon 2019; Woodworth, Girskis, and Walsh 2017). In cancer, these lineage maps can be used to understand how transcriptional programs evolve to overcome chemotherapy or seed metastatic sites. New techniques and data sets have spurred the development of a number of analysis tools, including tests for both gene and gene-set enrichment over the entire tree (Quinn et al. 2021; Schiffman et al. 2024; Forrow and Schiebinger 2021; DeTomaso and Yosef 2021; Lange et al. 2024; Price et al. 2022). Yet there is still a strong need for frameworks that can disentangle the contributions of genes to specific phenotypic endpoints, such as organ-specific metastasis or drug resistance, from those arising due to stochastic noise versus adaptive evolutionary pressures.

The field of phylogenetics has spent decades developing methods to understand evolution at the species level. Popularized by Butler and King in 2004, the Ornstein-Uhlenbeck (OU) model has become a valuable tool that captures both the forces of selection and random genetic drift in the evolution of a continuous trait (Butler and King 2004; Beaulieu et al. 2012). OU models describe patterns of variance of a continuous trait in a population with a shared history. Given a phylogenetic tree with a trait value per leaf, the OU model is used to test for different ‘adaptive landscapes’ (Methods). These can be thought of as hypothesized environmental niches, also referred to as *regimes*, each with a specific optimal trait value (θ). The model is further parameterized by the selection coefficient, ɑ, which describes how quickly a trait approaches its optimal value, and σ, which captures the rate of stochastic evolution by describing the variance of the Brownian motion component of the model, σ^2^ (Methods).

Here, we developed SCOUT, a quantitative framework for modeling gene expression evolution in single-cell lineage trees (**Figure 1A**). First, a set of evolutionary hypotheses is defined for a clonal population. We categorize evolutionary hypotheses into three groups using the definitions established by Hirsch et al. 2025: a *neutral*, Brownian motion model of evolution driven exclusively by genetic drift (BM1); a *constrained* model where all individuals in a tree are pulled towards a single optimal value (OU1); or an *adaptive* model where all individuals within the same niche, or regime, are pulled towards regime-specific optimal values (OUX, x = the number of regimes) (**Figure 1B**). For example, in a cancer setting, each metastatic tissue niche may present unique challenges to survival, which drives gene expression to different optima. The landscape of different optima thus outlines an *adaptive* evolutionary hypothesis. For each gene of interest, we fit the different hypotheses: neutral (BM1), constrained (OU1), and adaptive (OUx). The best-fit model per gene is selected, and the final output is a per-gene classification with accompanying parameter estimates for selection and drift.

**Figure 1.**
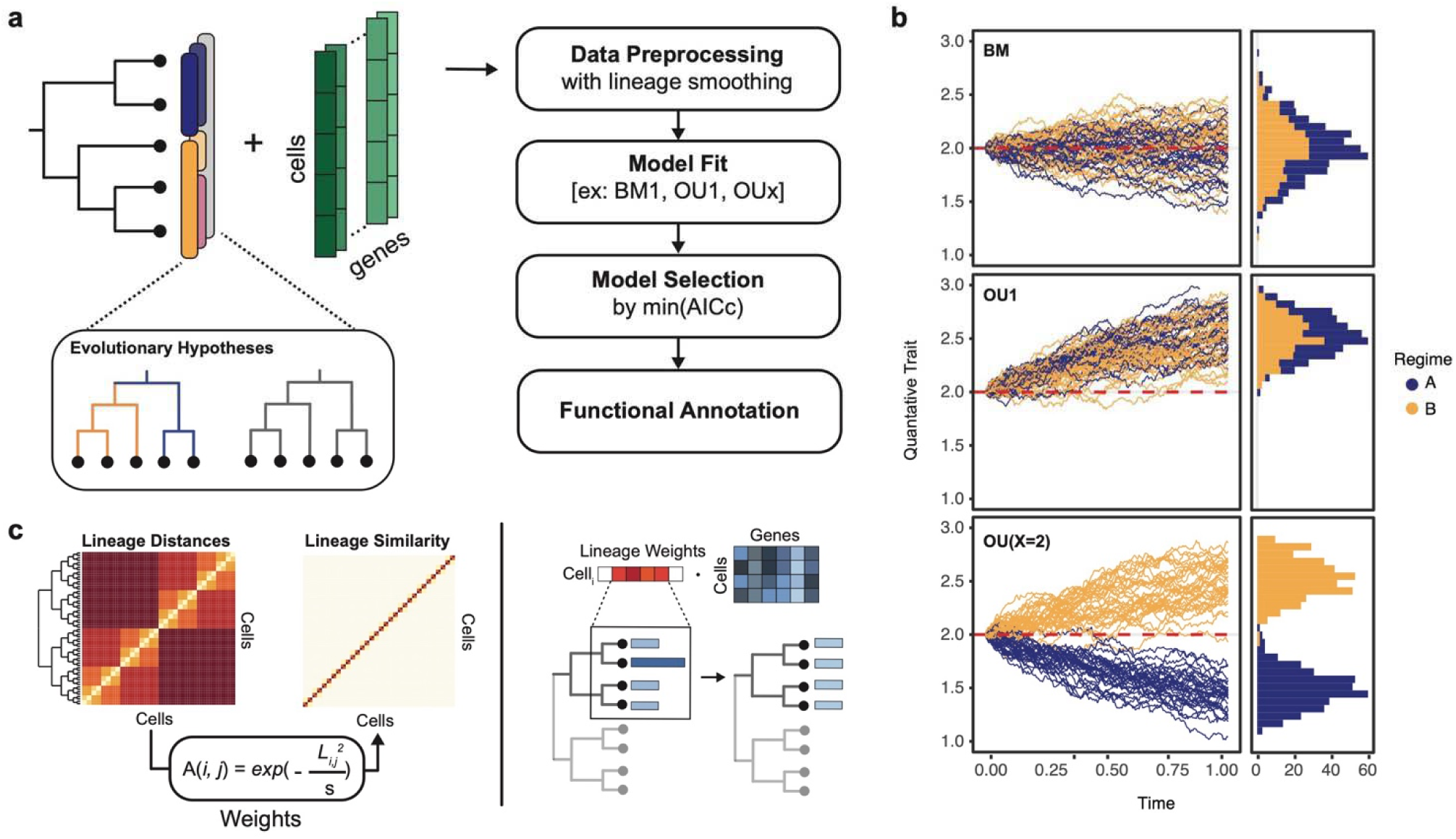
Overview of SCOUT. (a) Workflow of SCOUT. Given a single-cell lineage tree, a gene-by-cell counts matrix, and a set of evolutionary hypotheses, identify the best-fit evolutionary model per gene. Example evolutionary hypotheses illustrate how cells are partitioned into ‘regimes’, or evolutionary niches. (b) Simulated evolutionary models for neutral (BM1), constrained (OU1), and adaptive (OUx, x=2) traits. Each line represents the evolutionary trajectory of a terminal node with α=0.75 for the OU1 and OU2 models, and σ=0.25 for all models. All start with an initial value of 2, the optimal value (θ) for OU1 is 3, and the optimal values for OU2 are 1 and 3. As the traits continue to evolve to time infinity, a histogram illustrates that the distributions approach normal, or multivariate normal. (c) Proposed workflow to perform lineage smoothing on gene expression data. First, cell-by-cell distances are established based on a single-cell lineage tree. These distances are converted to similarities via an adapted Gaussian kernel. Lineage weights are restricted to the K lineage neighbors; cells further than the Kth on the tree are not considered in the smoothing for cell_i_. Log-normalized gene expression values are smoothed by taking the dot product of the lineage similarity matrix and the input gene expression. Smoothing parameter, k, describes the kth neighbor from the cell of interest. It is used to determine the maximum distance a cell can be from cell_i_, therefore establishing the width of the window (i.e. how many lineage neighbors to give weights to) and this maximum distance is also used as the bandwidth parameter, s.

Previous studies have explored the profiling of gene expression using an OU model (Hirsch et al. 2025; Chen et al. 2019). One recent study sought to profile subclonal cancer evolution by modeling gene expression as an OU process, utilizing single-cell gene expression and a lineage tree generated at the subclone level with 23 tips, limiting overall power (Cooper et al. 2016; Hirsch et al. 2025). Another study employed the OU process to model bulk gene expression across seven tissues in 17 species, providing evidence of stabilizing selection across species, albeit with limited power due to the species count (Price et al. 2022; Chen et al. 2019). Furthermore, a study more generally explored the theoretical basis for using OU models to model gene expression. The publication demonstrated that using bulk RNA-sequencing data gene expression counts may be biased due to unbalanced sampling of cell types and tissue composition. Thus, often these measurements do not capture the true variation in a gene’s expression when comparing between species (Price et al. 2022). These studies demonstrate both the potential of modeling gene expression as an OU process, but also key issues attributable to low-resolution data.

We hypothesize that the added power of single-cell lineage trees (often with 1000+ leaves), paired with single-cell RNA-sequencing, will enable us to accurately identify a gene’s adaptive landscape and estimate its selective constraints (Price et al. 2022; Cooper et al. 2016; Grabowski et al. 2023; L. S. T. Ho and Ané 2014). We first test SCOUT in simulated data with known adaptive landscapes and realistic single-cell gene expression profiles. Here, we introduce a lineage-based neighborhood smoothing approach to overcome the noise of single-cell data (**Figure 1C**). We then turn to a common gold standard dataset, the Caenorhabditis elegans developmental system, where single-cell transcriptomics data can be coupled to known cell lineage. Finally, we profile the genetic drivers of clonal evolution and metastasis using SCOUT, showing selection for both tissue-specific expression and microenvironment-interaction genes in a lung adenocarcinoma xenograft model with CRISPR/Cas9 insertion-deletion barcodes.

## Results

### Benchmarking in simulated data

We first sought to benchmark the use of Ornstein-Uhlenbeck models in scLT trees through simulation. This is particularly important given the sensitivity of OU models to measurement error and the well-known noise of gene expression (Price et al. 2022). Our goal was to generate biologically plausible single-cell counts data with realistic levels of noise and variation that reflect an underlying evolutionary regime structure. We first simulated single-cell lineage information, including final state assignments, based on a known cell-state transition tree with *TedSim (Pan, Li, and Zhang 2022)*. *TedSim* produces cell state annotations and a ground truth tree, as well as a character matrix recapitulating results seen with typical indel-based lineage barcodes (**Figure 2A**). Based on the cell states of the simulated population, we then generate realistic gene expression counts with known underlying evolutionary models. To do this, we generated typical OU traits with known α,σ and θ parameters. These values were then used to model extrinsic levels of variation in a single-cell population and infer per-cell and per-gene kinetic parameters, which were subsequently used to generate counts for a given cell and gene via a Beta-Poisson model (Methods) (Zhang, Xu, and Yosef 2019). This approach maintains the extrinsic variation among cells with respect to the lineage tree and known adaptive regime while also incorporating realistic levels of noise due to the transcription process.

**Figure 2.**
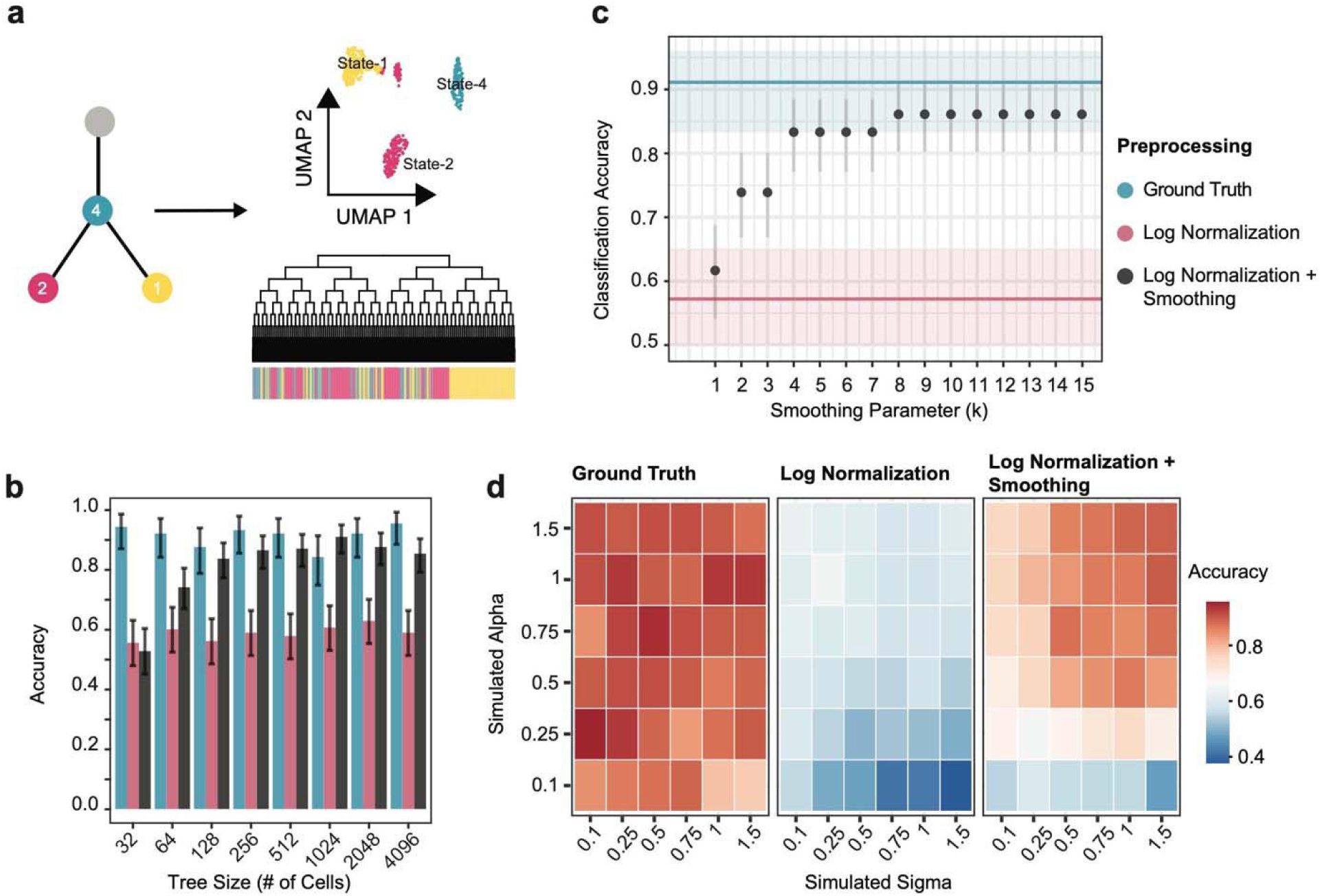
**Benchmarking SCOUT in simulated datasets.** (a) Schematic of simulation strategy. Given a known cell state tree with two terminal states and one intermediate state, TedSim returns a full tree with state labels and gene expression counts. We simulated additional genes with known underlying evolutionary models using a separate workflow (Methods). (b) Classification accuracy of SCOUT across tree sizes of 2^*g*^ for g=4,..,10. Simulated count matrices were preprocessed using standard log normalization and the log normalization with an additional smoothing step to minimize noise. These were compared against ground truth, which are the ‘OU’ continuous traits prior to transformation into counts (see Methods). For all trees and datasets, *α* = 0.75, σ = 0.75 and *k* = 8. Error bars represent a 95% confidence interval around the accuracy. (c) Classification accuracy of SCOUT across different smoothing parameters. The ground truth and standard log normalization accuracy results are displayed as horizontal lines with a 95% confidence interval in the surrounding shaded boxes. The counts were generated from an OU distribution simulated with parameters: *α* = 0.75, σ =0.75, and *N cells* =512. Error bars represent a 95% confidence interval around the accuracy. (d) Heatmap of accuracy results across a parameter grid search of α and σ, compared across the different preprocessing approaches.

As noted previously, OU models are sensitive to noise, which inflates the variation of a trait between species, or tips of the tree. This in turn reduces phylogenetic signal, the tendency of species that are more closely related to appear more similar to each other, and can erroneously favor a constrained model over a neutral model (Grabowski et al. 2023; Cooper et al. 2016) (**Figure S1A**). Our single-cell smoothing approach aims to restore the phylogenetic signal that has been reduced by noise. Briefly, we leverage lineage distance to assign weights to a cell’s most closely related neighbors via an adapted Gaussian kernel (methods). We then smooth log-normalized gene expression counts by taking the dot product of the lineage similarity matrix and the normalized counts. Smoothing is parameterized by a single parameter, k, which identifies the ‘kth’ farthest lineage neighbor from a cell of interest. It is used to both identify the width of the neighborhood and the bandwidth, s, of the kernel **(Figure 1C)**. Similar approaches have been implemented to smooth or impute gene expression values using spatial coordinates or transcriptomics (van Dijk et al. 2018; Holdener and De Vlaminck 2025) (Methods).

We set out to benchmark SCOUT in our scLT experimental setting. Our first priority was to determine how well we could classify genes based on their true underlying model in a single-cell setting using different preprocessing strategies. We simulated datasets across different-sized trees, ranging from 32 to 4096 terminal single-cells. For each tree, we generated 50 genes for each of three models: the neutral model (BM1), the single-regime constrained model (OU1), and the multi-regime adaptive model (OU3). For the multi-regime adaptive model, regimes were assigned based on the simulated cell state per *TedSim* as previously described.

As expected based on previous literature, the 32-leaf tree resulted in the worst performance with an overall classification accuracy of 0.55 [95% CI: 0.47, 0.62] using standard log normalization and 0.52 [95% CI: 0.44, 0.59] with the additional smoothing step (K = 8 for all trees) (**Figure 2B**) (Chen et al. 2019; Price et al. 2022; Cooper et al. 2016). Across the remaining tree sizes, preprocessing the data with standard log normalization results in consistently low classification accuracy. In contrast, the smoothing step greatly improves classification accuracy, achieving accuracy that approaches the ground truth for trees as small as 128 leaves. Furthermore, we observed that smoothing reduces the misclassification of true neutral genes as constrained, relative to log-transformation alone (**Figure S1A**). Finally, as the smoothing parameter, k, increases, we observe that the classification accuracy in our smoothed dataset rises to meet the ground truth results, representing a large increase over log-normalized data (**Figure 2C**).

Next, we performed a parameter grid search using our simulation framework to explore which ranges of the selection parameter, α, and the genetic drift parameter, σ, can be reliably recovered from scLT data. Across parameter combinations, we observe a consistent improvement in performance with increasing tree size in smoothed data, with higher values of ɑ and σ, resulting in higher accuracy. Whereas with log-transformation alone, accuracy improves nominally with increasing tree size (**Figure S1B, Supplemental Table S1)**. For both the log-normalization and smoothing preprocessing strategies, performance was poorest when σ was high and α was low. Under these parameters, the process becomes nearly indistinguishable from Brownian motion, especially at very low α values. This matches intuition: as α decreases, the model approaches neutrality, increasing the likelihood of misclassification. Accuracy again improved substantially when the data were smoothed rather than only log-transformed, across a broad range of parameter combinations (**Figure 2D**). Taken together, our simulations demonstrate that SCOUT can accurately classify the correct evolutionary model under realistic conditions for single-cell lineage trees.

### C. elegans

We next sought to evaluate SCOUT in a real-world lineage dataset. The *C. elegans* lineage tree is fully resolved, and a single-cell atlas has been mapped to the worm’s lineage tree (Sulston et al. 1983; Packer et al. 2019). These data are frequently used to benchmark methods that analyze single-cell lineage tracing data, as they provide a ground truth setting for scLT experiments (Lange et al. 2024; K. Wang et al. 2024; Forrow and Schiebinger 2021). Here, we used the known lineage tree and single-cell gene expression data to understand the adaptive landscape driving development of the nematode worm.

In the Packer dataset, multiple embryos were sequenced, and many single cells were sampled from the same point on the lineage tree, including intermediate lineage states. To more closely mirror the data from a scLT experiment, we only considered cells with a terminal (leaf) lineage state (5,792 cells; 391 terminal states). To generate a test case, we randomly sampled one cell per terminal state to achieve a ‘pseudoembryo’ of 391 cells (K. Wang et al. 2024). For genes expressed in at least 20% of cells (N=3819), we fitted the following adaptive landscape models: [BM1, OU1, OU2, and OU6]. The OU2 regimes are determined by the AB versus P1 lineage split, whereas the OU6 regimes are defined by the major lineage branch of the cell (AB, MS, C, D, E, Z). These two adaptive hypotheses were based on the tendency for cells to maintain their relative position, or niche, in the worm throughout development (**Figure 3A**) (J. Liu and Murray 2023). These lineage assignments unite cells originating from different UMAP clusters in the transcriptional space (**Figure 3B-D**).

**Figure 3.**
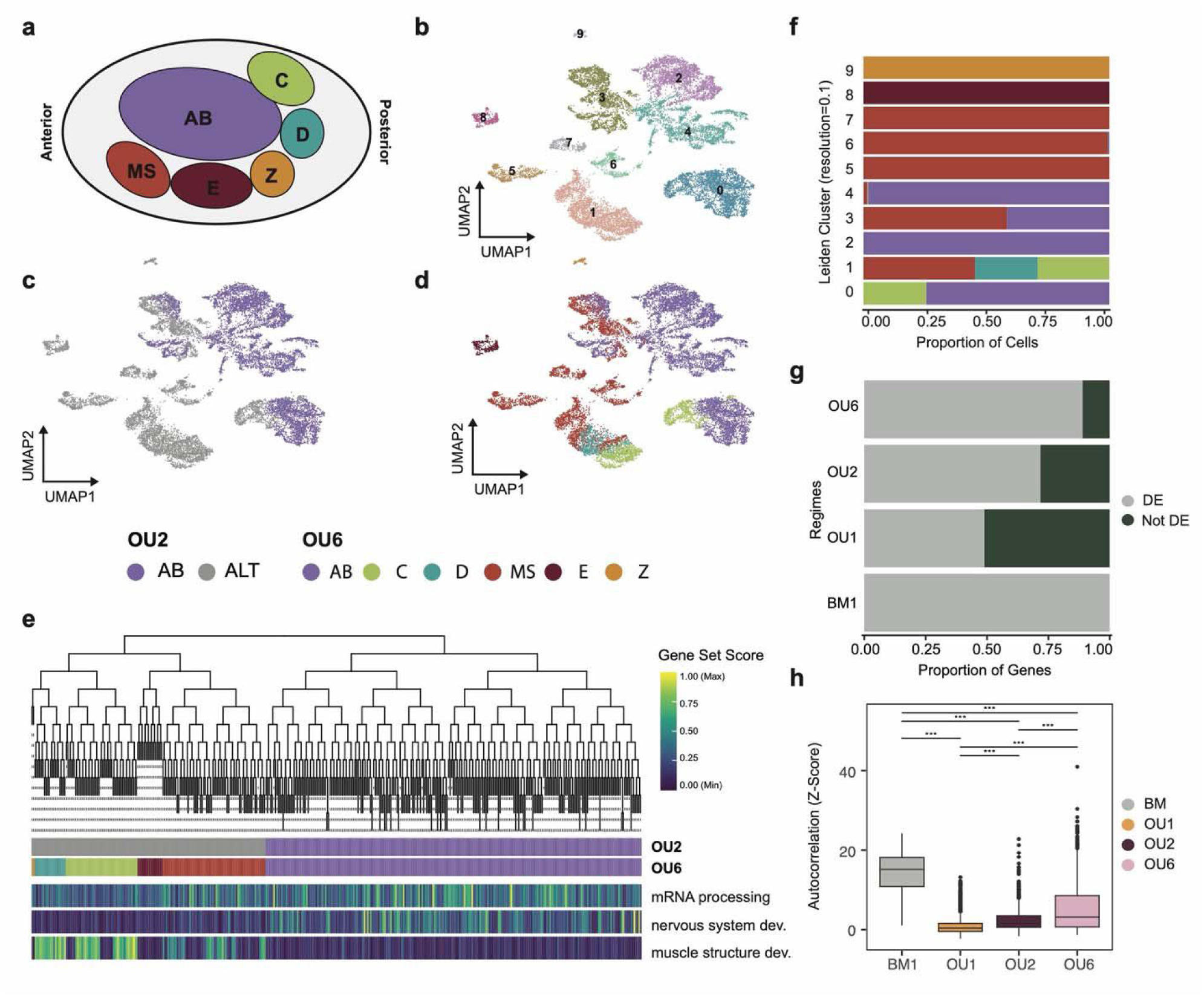
Detecting selection dynamics in *C. elegans* development. (a) Approximate representation of C. elegans founder cells positioning in the developing embryo. (b-d) UMAP of *C. elegans* terminal state cells (Number of cells = 15,792, number of terminal states = 391) colored by membership to adaptive regimes (b) colored by cluster membership (resolution = 0.1), (c) OU2, AB lineage versus P1 (not AB) and (d) OU6, the major lineage. (e) *C. elegans* lineage tree where each leaf represents a unique terminal state. Color bars denote the two adaptive evolutionary hypotheses being tested. Heatmap representation of GO terms along the cells in the tree. GO term scores were calculated using scanpy’s ‘score_genes’ function based on the genes overlapping with the term. (f) Normalized bar plot outlining the per-lineage-branch (based on an OU6) breakdown within Leiden clusters. (g) Normalized bar plot of genes assigned to each class of model, split by whether the gene was detected in differential gene expression analysis based on Leiden clusters. (h) PATH auto-correlation Z-score results. Genes are colored by their model classification. Statistical significance was determined by performing a Wilcoxon test with significance levels: p-value < 0.001: ***, p-value <= 0.01: **, p-value <= 0.05: *, p-value > 0.05: NS.

Ultimately, one-third of genes tested could be confidently assigned to a final model. Unassigned genes were removed because their AICc improvement over the next-best model was insufficient (ΔAICc < 2). The majority of these unassigned genes were OU1, of which 96% had OU2 as the next best model, suggesting that it can be challenging to distinguish between two different, non-Brownian-motion models. Of the genes classified, 30 were assigned to a Brownian motion model; the remaining 1238 were assigned to one of the adaptive evolution regimes. This is consistent with our expectation that the majority of genes will be under constrained transcription (selection) in development. 519 genes were identified as OU1, 217 genes were identified as OU2, and 502 genes followed an OU6 model (**Supplemental Table S2**).

To evaluate the biological plausibility of SCOUT classifications, we performed an over-representation analysis of the assigned genes. Genes under an OU1 were enriched for general GO terms involved with mRNA processing or protein transport, fundamental cellular processes that would be expected to be under selection towards a common optimal value. For the adaptive genes, we first performed hierarchical clustering on the optimal expression value (θ) for the OU6 and OU2 genes to ensure we captured genes with common optimal value patterns (**Figure S2A, B**). Those genes assigned as OU2 with a higher optimal value in the AB lineage were enriched for GO terms such as neurogenesis and neuron development, which is consistent with the fact that the AB lineage is the largest producer of neurons (Sulston et al. 1983) (**Figure 3E, Figure S2 C,D**). Finally, genes found under an OU6 regime were enriched for highly specific GO terms such as muscle differentiation in the C and D lineages, containing known lineage branch points for body wall and other muscle populations (**Figure 3E, Figure S2 E, F; Supplemental Table S3**) (J. Liu and Murray 2023).

Next, we were curious how SCOUT’s classifications compared with differential gene expression results based on Leiden clustering across all cells with a terminal lineage annotation. The single-cell clustering produced 10 clusters, with some clusters restricted to specific lineage groups such as Z and E (exclusive to clusters 9 and 8, respectively) and others, such as clusters 0, 1, and 3, split across multiple lineage clades (**Figure 3B, F**). We performed differential gene expression to identify enriched genes in each cluster and filtered for FDR=0.1 and log-fold change > 1.5. The resulting gene list was then compared to the OU classifications. Notably, genes following the OU6 model showed the highest overlap with the DEG list, whereas OU1 genes showed the lowest overlap (**Figure 3G**). As OU1 genes have a unified distribution, they would not be expected to be detectable by differential gene expression. While we do see limited OU1 genes in the DEG list, this is likely due to a very permissive DEG list, which includes any gene that passes the filter for any cluster.

Finally, we tested our SCOUT-classified genes for phylogenetic signal. Recent work has explored the use of Moran’s I, a statistic that describes spatial autocorrelation, to profile heritability in single-cell lineage trees (Schiffman et al. 2024). Genes that are heritable have high autocorrelation Z-scores, indicating that a neighbor is likely to have a similar level of expression for the same gene. Phylogenetic signal is notably distinct from the study of evolutionary processes (Revell, Harmon, and Collar 2008). However, it is frequently reported that traits evolving under Brownian motion exhibit high phylogenetic signal, as the variation within the trait is purely dependent on time, embodied here by the distance between species on the tree. We observe that genes found to evolve under an OU1 process had the lowest autocorrelation Z-scores, with the majority having Z-scores below 2. OU2 and OU6 genes had significantly higher autocorrelation Z-scores. (**Figure 3H**). As expected, genes identified as being under a BM model exhibited the highest autocorrelation Z-scores. These results suggest that SCOUT produces reasonable gene classifications that behave as expected using ‘real’ single-cell gene expression data.

### LUAD Metastasis

Tumor cells migrate from primary tumor sites and adapt to a new environmental niche at distant tissue sites. To explore this biological process from an evolutionary perspective, we applied SCOUT to CRISPR single-cell lineage trees from a lung adenocarcinoma (LUAD) xenograft model (**Figure 4A**) (Quinn et al. 2021). We evaluated 68 trees from the original publication with SCOUT, testing each gene for neutral evolution under a BM1 model, constrained evolution under an OU1 model, and adaptive evolution under an OU4 model (**Supplemental Table S4**). For the adaptive OU4 model, we assigned regimes to the primary tumor in the left lung (LL) and metastatic sites in the right lung (RL), mediastinum (M), and liver (Liv). Across all trees and genes, the top model improved upon the next best by an average delta AICc of 7.72 [95% CI: 7.54, 7.89]. An average of 924 (∼30%) genes were removed due to ambiguous model assignments. Tree size was correlated with average delta AICc (r = 0.604, p-value < 6.351e-08) and AICc weights of the top model were significantly higher in genes with a delta AICc > 2 (**Figure S4A, B**). Across all trees, the majority of genes fell under a simple OU1 model of constrained evolution (**Figure 4B**). We observed a moderate but significant negative correlation between the proportion of genes annotated as neutral (BM1) and tree size (r = -0.615; p-value = 4.3e-06 for N=47 trees with at least one Brownian motion gene) (**Figure 4B**).

**Figure 4.**
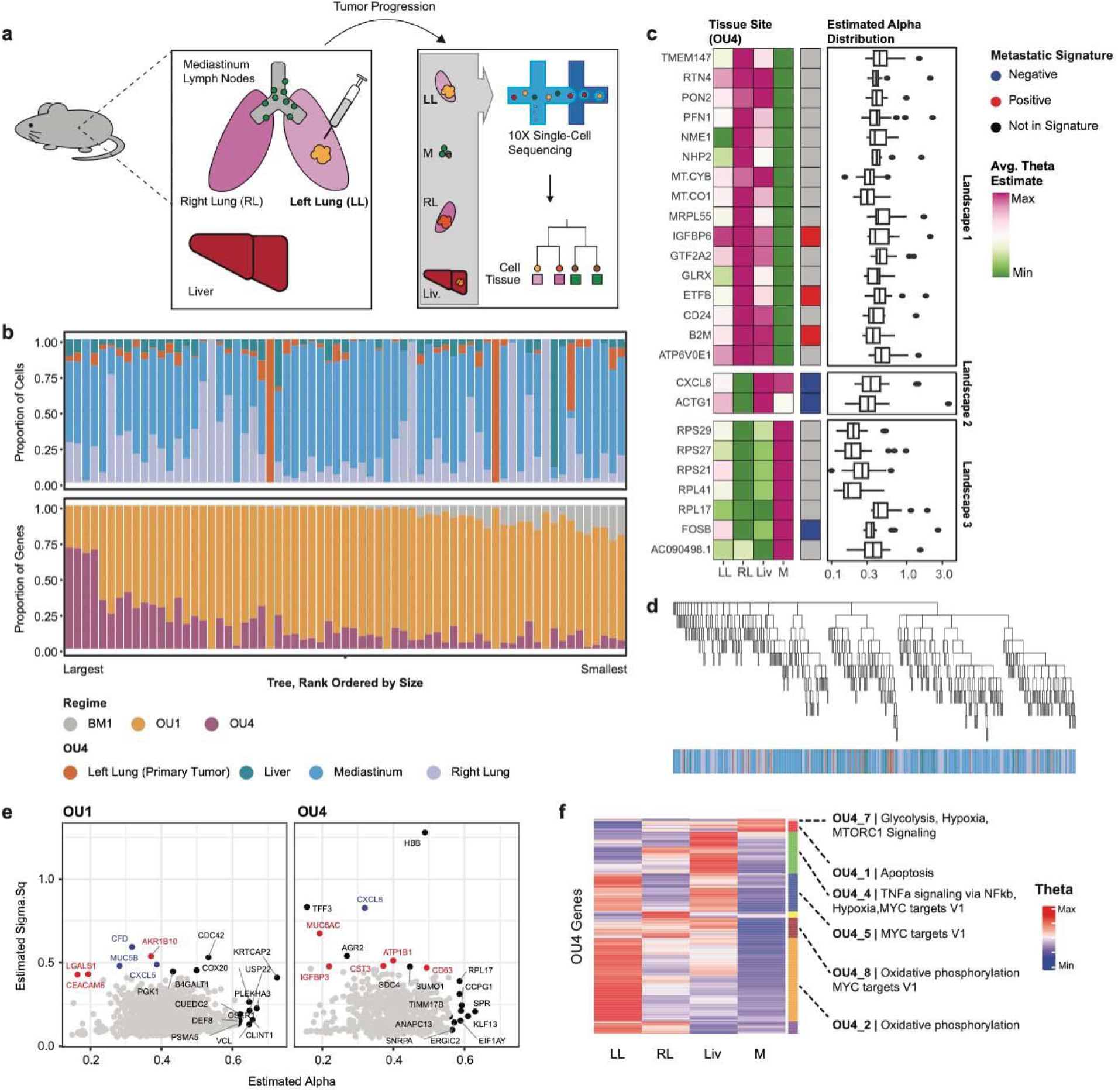
**Detecting selection dynamics in lung adenocarcinoma metastasis.** (a) Experimental design of Quinn et al. 2021 dataset. A mouse was implanted with an engineered A549 cell line, equipped with lineage tracing capabilities. The cancer was allowed to metastasize, and cells were taken from four metastatic sites. These samples underwent single-cell RNA-sequencing to extract cell state and lineage information simultaneously. (b) Normalized bar plot of per-cell tissue composition (upper) and per-gene model classification (lower) per tree. Tissue composition is defined through the lens of an OU4. Trees are approximately ordered by size from largest to smallest after pruning for cells with both lineage and transcriptomic information. (c) The top 25 common genes annotated as an OU4 across all trees tested. Boxplots represent the distribution of alpha values per gene across trees with the shared annotation. A min/max scaled heatmap illustrates the average theta (optimal) value across trees with the same annotation for that gene. Genes are split into “patterns” that have similar adaptive landscapes based on their optimal value profiles. The middle heatmap denotes whether the gene overlaps with the metastatic signature as defined in the original publication. (d) Dendrogram representation of LG13 tree with OU4 regime annotation heatmap. (e) Scatter plot of constrained and adaptive genes for LG13 by their alpha and sigma parameter estimates. Genes in the top 10 alpha or sigma-squared values are annotated by name and colored by their inclusion, or not, in the metastatic signature. (f) Heatmap of min/max normalized theta values per gene annotated at an OU4 gene for LG13. Genes were clustered to identify groups of genes with similar evolutionary structure across the primary and metastatic sites. Clusters are annotated along the rows. Up to the top 3 HALLMARK gene signatures per cluster are annotated.

We were first interested in which genes were consistently favored by one model of evolution in multiple trees. Genes relevant to tumor progression in LUAD, such as GOLPH3, MAFK, AGRN, and CD164 were labeled as OU1 in at least 80% of trees. To explore consistently-annotated OU4 genes, we compiled a list of the top 25 adaptive genes by number of trees. On average, these top genes were found in 28 trees [95% CI: 26, 29]. We observed that some of these genes shared similar adaptive landscape profiles. To better delineate common landscapes, we clustered genes based on the average optimal value for each metastatic site for all trees with that gene-model combination. Genes in landscape 1 were shared by the LL and M, having the highest values, and RL and Liv had lower to moderate values. Among these genes are some cross-referenced with the Quinn et al metastatic signature, such as IGFBP6 and B2M. Landscape 3 is in opposition to pattern 1, featuring high RL and Liv optimal values, moderate LL optimal values and low M optimal values. Genes in this landscape are largely ribosomal proteins with the exception of FOSB. Landscape 2 exhibits low optimal values in RL and M sites, but high values in LL and Liv sites. Landscape 2 includes only two genes, CXCL8 and ACTG1. CXCL8 is known for its role in modulating the tumor microenvironment (TME) and is annotated consistently as OU4 across 34 trees. ACTG1 is also known for its role in increasing metastatic potential (**Figure 4C**) (Suresh and Diaz 2021). These results align with our expectations that OU1 genes would be globally selected while OU4 genes demonstrate more clone, and tissue-specific patterns.

We next turned to a single tree, LG13 (N=425 cells), a highly metastatic clonal tree highlighted in the original publication (**Figure 4D**). We hypothesized that SCOUT could detect genes under meaningful selection, providing insight into the clone’s highly metastatic behavior. In LG13, we tested 3164 genes found in at least 20% of cells. Of genes confidently assigned (N=2340), SCOUT identified 26.7% of genes as being under an OU4 model and the remainder under an OU1 model. We again observe that genes annotated as OU4 have significantly higher autocorrelation Z-scores than OU1 genes **(Figure S5B).** To zoom in further, we prioritized genes based on parameter estimates, split by evolutionary model classification **(Figure 4E)**. Interestingly, the genes listed in Quinn et al.’s metastatic signature feature high sigma values. A number of these high-sigma genes (CXCL5, CXCL8, AGR2) across both adaptive and constrained models are associated with modifying the tumor microenvironment to support tumor progression (Q. Liu et al. 2016; Xiong et al. 2022; Sun et al. 2024; Fessart et al. 2021; Huang et al. 2014). 𝝈^2^ is often interpreted as the stochastic evolutionary rate, describing the variance in a trait after a certain period of time. This suggests TME genes’ expression are subject to a high rate of stochastic evolution, potentially equipping cells to be more adaptable to new tissue environments. We identified the high-alpha genes USP22 and PSMA5 for their involvement in tumor progression via PD-1 upregulation, which enables immune evasion in cancer cells (Hu et al. 2015; Lu et al. 2022; Yao et al. 2025). The adaptive (OU4) genes, such as MUC5AC and IGFBP3, promote metastasis and invasion, whereas the constrained genes (OU1) are directed at immune evasion (Ponnusamy et al. 2013; Chaudhary et al. 2024; Y. A. Wang et al. 2017).

We next performed differential gene expression analysis on cells from the LG13 tree. When performed using tissue sites to group cells, only 7 genes survive filtering with cutoffs FDR=0.01 and log fold-change = 1.5. Yet of these 7 genes, 5 overlap with LG13 OU4 genes. All 7 genes were upregulated in the liver metastatic site (Liv), except for HBB, which was upregulated in the right lung (RL). We then performed Leiden clustering on the LG13 cells and identified four clusters. We reran the analysis and recovered 43 differentially expressed genes that passed filters, with all clusters having at least one upregulated gene. Approximately 67% of genes overlap with either OU1 or OU4 gene lists. The OU4 genes overlap the DEG list (Jaccard similarity = 0.026) more than the OU1 genes (Jaccard similarity = 0.007). Of the top five enriched genes per cluster, CXCL8 and CXCL family genes are enriched in cluster 3. Interestingly, no cells from the primary site (LL) are present in cluster 3 **(Figure S5 C)**. This aligns with the expectations given the optimal value profile of CXCL8 across 34 trees, which shows low values of CXCL8 in the lung sites and high values in Liv and M sites **(Figure 4C)**. TFF3, upregulated in cluster 0, is also annotated as OU4 with a high sigma value in LG13 **(Figure S5 D)**.

Finally, we sought to deconvolve the OU4 genes into common adaptive landscapes. We used hierarchical clustering to partition genes with similar optimal expression patterns, identifying eight unique landscapes that varied in size from 7 to 243 genes. To contextualize genes within the different landscapes, we performed an overrepresentation analysis on the genes in each group using the HALLMARK gene sets. We found enrichment of HALLMARK gene sets in 6 out of 8 landscapes (**Figure 4F; Supplemental Table S5**). The largest group describes genes involved in oxidative phosphorylation and has the highest optimal value in the primary tissue site, LL. The smallest group, with 7 genes, featured genes involved in hypoxia, glycolysis, and the MTORC1 signaling pathway. This suggests an activation of pathways needed to provide cancer cells with the energy to survive in a new environmental niche. This small set of genes is expressed highly in the mediastinum and lower in the RL and Liv tissue sites. Landscape 4 co-opts TNFa signaling via TNkb, hypoxia, and MYC targets. This highlights an additional trajectory that fuels this clone’s aggressive metastatic identity. Together, these results demonstrate that SCOUT can capture key transcriptional programs underlying tumor progression, which exhibit unique selective pressures across different metastatic sites, and point to the survival avenues taken by individual clones to adapt to new environments.

## Discussion

Cellular plasticity allows cells to respond to their environment in both normal development and in disease. This plasticity acts at multiple time scales, preserving transcriptional programs already well-suited to their environment, rerouting others to respond to transient challenges, or to adapt to new stimuli. The challenge is then to disentangle complex cellular translational programs into these different adaptive regimes. Single-cell lineage tracing provides the framework to capture this plasticity, using the tools of phylogenetics to address these fundamental questions.

Here we present our method SCOUT, which leverages an Ornstein-Uhlenbeck (OU) model with single-cell data smoothing to dissect gene expression evolution in lineage trees. We first benchmarked SCOUT using simulated single-cell lineage data to understand how the model performs in the presence of scRNA-seq data noise and to determine the extent to which evolutionary models can fit to the data. Our simulations yielded promising results, accurately identifying the correct model with reasonable parameter estimates. Using *C. elegans* as a validation dataset, we next identified gene classifications based on cell lineage assignments, which provide an orthogonal view of genes outside of canonical differential gene expression analysis. Finally, we demonstrate compelling biological processes and genes under different regime structures across highly versus weakly metastatic clones in lung adenocarcinoma.

There are limitations with our approach. SCOUT relies upon local smoothing of the single-cell transcriptional space to overcome stochastic noise and measurement error. This is currently implemented as a hyperparameter; however, more adaptive approaches could be utilized in the future. Although single-cell lineage trees benefit from a large number of samples and we’ve chosen a relatively conservative model selection approach (ΔAICc > 2), measurement error could still bias our results (Silvestro et al. 2015). Larger trees will also require more computational resources, and some of the largest trees in the cancer dataset were downsampled for computational efficiency. Lastly, the trees analyzed in the paper are the product of imperfect technologies; all single-cell lineage tracing analysis methods will benefit from advances in the underlying technology to create higher-resolution and more accurate trees.

In conclusion, SCOUT establishes a framework for modeling the evolutionary dynamics of gene expression in single-cell lineage trees. SCOUT unites lineage information with phylogenetic modeling, providing a quantitative means to separate neutral, constrained, and adaptive processes that shape cellular behavior. As single-cell lineage tracing technologies continue to mature, approaches such as SCOUT will enable increasingly precise views of how selective pressures and stochastic variation together sculpt developmental trajectories and disease evolution.

## Methods

### SCOUT

#### Overview

SCOUT models gene expression as the Ornstein-Uhlenbeck process to detect selection using single-cell lineage trees. The OU model has three parameters: alpha (α), a measure of the strength of selection on a continuous trait; theta (θ), which identifies the optimal value of the trait; and sigma (σ), such that σ^2^ is the rate of stochastic evolution and controls the variance of the genetic drift component of the model. The model describes the change in a continuous trait *X_i_* over an increment of time. As time approaches infinity, the trait distribution converges to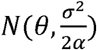. Notably, as α approaches 0, the model resembles pure neural drift.

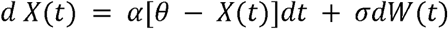

The ultimate value of the OU model is its ability to test against other evolutionary hypotheses to identify the best-fit adaptive landscape for a continuous trait (Butler and King 2004). As input, SCOUT accepts a single-cell lineage tree in Newick format and a metadata file containing evolutionary hypotheses, species ID (the cell barcode, or leaf label in the tree), and the continuous trait(s) of interest (genes). Genes can be pre-normalized or raw counts. In the absence of branch lengths, for simplicity, all edges are assigned a length of 1.

We use the *OUwie* implementation of the OU model (Beaulieu et al. 2012). To perform likelihood calculations, we use the ‘three.point’ algorithm. Briefly, the ‘three.point’ algorithm improves computation time by avoiding calculating the full matrix inverse of the phylogenetic variance-covariance matrix as described in detail in previous work (L. si T. Ho and Ané 2014). *Ouwie* also allows for the use of the original ‘inverse’ method which is more computationally intensive and therefore not recommended for large trees. Otherwise, *OUwie* is run per gene-and-model with default parameters.

#### Ancestral Character Estimation and Defining the Adaptive Landscape

An essential element of SCOUT and more generally, the use of OU processes to model evolution, is defining a set of evolutionary hypotheses, or adaptive landscapes. An evolutionary hypothesis describes a partitioning of leaves into different environmental niches, or regimes, that informs how a trait will evolve. In a neutral or constrained model of evolution, all leaves are within the same ‘regime’. But in an adaptive model, leaves are assigned to different ‘regimes’. Once assigned, ancestral character estimation is performed to annotate the internal nodes with a regime using the *ape::ace* implementation for discrete characters and an equal rates state transition model (Beaulieu et al. 2012; Paradis, Claude, and Strimmer 2004). This method requires trees to be fully dichotomous (number of nodes = number of tips - 1). Final regime assignments for internal nodes are made by taking the state with the maximum likelihood.

#### Model Selection

To pick the best fit model per gene, we select the model with the minimum AICc. For datasets that require an extra level of stringency, we keep only gene-model combinations with a delta AICc to the next best model of greater than 2. Finally, based on our simulations, for ‘real’ datasets we removed constrained and adaptive genes that had very small alpha values (<0.1) because these are practically indistinguishable from a neutral model.

#### Smoothing

To address the issue of noise introduced by using single-cell gene expression data, and the sensitivity of OU models to measurement error, we introduce a smoothing step during preprocessing. We smooth gene expression values based on the assumption that cells that are closer to each other in a tree will be more similar, essentially aiming to recover phylogenetic signal lost to noise. We take an approach that aims to preserve local structure as much as possible. We based our strategy on similar approaches implemented in spatial and transcriptomics datasets, which incorporate flexible bandwidth parameter estimates depending on the density of the distance matrix and only consider neighbors within a certain window from the cell of interest (van Dijk et al. 2018; Holdener and De Vlaminck 2025). Here, given a single-cell lineage tree, we can obtain a cell-by-cell lineage distance matrix based on the tree structure. We then convert the distances to similarity weights using an adapted Gaussian kernel strategy. For lineage distance matrix, *L* lineage similarity matrix, *A* and cells *i* and*j*:

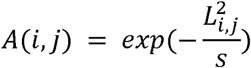

We define bandwidth parameter, *s*, per cell such that *s* = *max*(1,*L*(*i,k*)), where *k* represents the *k^th^* farthest cell from *cell_i_*. We further limit the smoothing to only the closest, local, cells by setting the similarity of all cells farther than the *k^th^* farthest cell to zero. The final window of weights is normalized row-wise so that all weights sum to 1 and the diagonal is set to 1 to ensure a cell is most weighted by itself (van Dijk et al. 2018). The final result is a local-weights matrix. This is then used to transform the log-normalized gene expression counts by taking the dot product of the log-normalized counts and the local weights to return a final smoothed cell by gene matrix.

### Simulation Design

#### Simulating lineage

To simulate gene expression and lineage data, we leverage two closely related tools: TedSim and SymSim, as well as the *OUwie.sim* function from the R package OUwie (Pan, Li, and Zhang 2022; Zhang, Xu, and Yosef 2019; Beaulieu et al. 2012). First, we use TedSim to simulate trees of a specific size based on a *cell state tree* which outlines the cell state differentiation process. Using this *cell state tree*, TedSim relies on the concept of asymmetric cell division to model cell state changes. To generate the dataset(s) in the manuscript, we use parameter estimates of probability of asymmetric division = 0.4, and step size = 0.4, to simulate cell state assignments along the lineage (Lange et al. 2024). We simulated lineage for character sites (N_char) = 64, which could equate to 8 target arrays of 8 editable sites and *cell state tree* = ‘((t3:1, t4:1):1);’. This tree produces a dataset with three states.In addition to a lineage tree and character matrix, TedSim also produces internal node state annotations.

#### Simulating OU-genes

Instead of using the TedSim gene expression counts, which have no basis in any specific OU process, and to maximize our control over the simulation, we customized the gene expression counts simulator pipeline, SymSim (Zhang, Xu, and Yosef 2019). Briefly, SymSim, aims to simulate gene expression counts incorporating three elements of variation fundamental to single-cell datasets: (1) Extrinsic variation can be akin to biological differences like cell type or state, (2) intrinsic variation captures the fundamental processes of transcription, and (3) technical variation which is the error introduced through experimental processing. SymSim models intrinsic variation using the kinetic model of transcription which is parameterized by *k_on_ promoter on rate), k_off_ (promoter off rate), and s (synthesis rate)*. Ultimately, counts are generated following a Beta-Poisson distribution such that *p* = *Beta (k_on_, k_off_)* and *X* = *pois(p*s)*. The parameters *k_on_, k_off_,* and *s* are generated per-gene-and-cell taking into account the distribution of cell states, which are continuously represented as a multi-variate normal distribution for a multi-state example. Yet, in the typical SymSim pipeline, these cell state assignments are lineage-agnostic. Therefore, in order to preserve the variance structure of a trait that is informed by a lineage tree, we interject an OU-informed trait distribution that is substituted for the multivariate normal random variable canonically used in the SymSim pipeline to model extrinsic variation. To generate the OU-trait distribution, we use the OUwie function *OUwie.sim* which accepts as input a tree with internal node state annotations, leaf regime/state assignments, and the parameters α, σ^2^, and θ. This returns a continuous vector of length N-tips with an appreciable phylogenetic variance-covariance structure. Using this with the SymSim pipeline produces a count matrix of genes by cells with appreciable evolutionary structure.

#### Simulating adaptive genes in 293Ts

To take a separate look at simulating realistic gene expression counts, we leveraged a publicly available 293Ts dataset from 10X (https://www.10xgenomics.com/datasets/293-t-cells-1-standard-1-1-0). From the filtered barcode matrix, we took the top 100 highly expressed genes by average expression. From these data, we randomly sampled 512 cells that will be the leaves in our single-cell lineage tree. To simulate cell states, we generated a pure-birth tree using *phytools::pbtree()* with type=’discrete’ and then *castor::simulate_mk_model()* to simulate the evolution of a discrete trait based on a tree and a transition matrix (Revell 2024; Louca and Doebeli 2018). We simulated a two state model. To mimic OU2 genes, we multiplied half the genes in one of the two states by a factor of 3 to shift the gene expression distribution for those cells.

#### Evaluation

Classification accuracy was evaluated using the *caret* package which defines accuracy as the proportion of correctly classified instances across all classes for a task with more than two classes (Kuhn 2008). 

### C.elegans

#### Dataset

We used the publicly available *C.elegans* dataset available through the package *moscot (Klein et al. 2025)*. This dataset is a subset of the original Packer et al. experiment, reduced to cells with a lineage annotation (Packer et al. 2019). A lineage state in the *C. elegans* dataset is described by the starting lineage (AB, MS, E, C, D, Z) and then subsequently by the directionality of the observed cell split (*a,* anterior; *p*, posterior; *l*, left; *r*, right). There are multiple ways of subsetting these data further. While all cells in this dataset have a lineage annotation, not all cells are assigned a unique annotation. Meaning, there are a number of potential lineages they could belong to. Cells that are uniquely mapped to a single lineage are in the ‘complete’ lineage subset. A separate, non-overlapping, set of cells belong to the ABpxp lineage which is symmetrical (such that x=r or l). The final subset strategy resolves ambiguity in the cells that could not be uniquely mapped by randomly selecting one of their possible lineage assignments. As such, this is often called the ‘random precise lineage’ (Forrow and Schiebinger 2021; Lange et al. 2024; Klein et al. 2025). While the first two subsets benefit from confident annotation, they suffer from few terminal cells (both under 50). Instead, we relied on the ‘random precise’ subset and generated a ‘pseudoembryo’ where one representative cell per lineage state is randomly selected. Of the 391 terminal lineage states, each was represented by 40 cells on average (median 33) (K. Wang et al. 2024).

#### Single Cell Analysis

Subsetting the original *moscot* c. elegans dataset to the 15K cells that have a terminal lineage annotation and a known cell-type, we then re-clustered the remaining cells with Leiden community detection algorithm in Scanpy using resolution = 0.1 to generate 10 clusters (Klein et al. 2025; Wolf, Angerer, and Theis 2018). We used the *rank_genes_groups* function with method=’wilcoxon’ to perform differential gene expression analysis on our Leiden clusters. We labelled any gene with an adjusted p.value <= 0.01 and abs(logFoldChange) > 1.5 in any cluster as being ‘differentially expressed’. This list of genes was then compared for presence or absence to the results of SCOUT per evolutionary model tested.

We conducted over-representation analysis via *clusterProfiler* R package using the *enrichR* function (Yu et al. 2012). For *C.elegans* we used the Gene Ontology Biological Process aspect with organism database *org.Ce.eg.db* and q.value/p.value cutoff = 0.05. To break the gene groups into more discrete functional groups, we took the adaptive genes for OU2 and OU6 and clustered genes to identify groups of genes with similar optimal values across the different regimes. Clustering was performed by hierarchical clustering using Euclidean distance and then partitioned into groups using cuttree. The OU2 genes were clustered into two groups and the OU6 genes were split into four groups (**Figure S2**).

#### PATH

To evaluate the levels of phylogenetic signal within our genes and our tree, we used a recently published tool, PATH (Schiffman et al. 2024). Briefly, PATH calculates a spatial autocorrelation coefficient, Moran’s I, which analyzes the spatial distribution of continuous data. High *Moran’s I* indicates that a neighbor is more likely to look like itself than different. In terms of gene expression, this equates to cells closer on a tree having similarly high levels of expression for a gene with high Moran’s I. This is interpreted as a measure of phylogenetic signal. We implement PATH using the same set of genes as is input for SCOUT. Genes counts were preprocessed via log-normalization. We obtain a phylogenetic weight matrix through *inv_tree_dist* which returns the inverse tree distance which is then used as input to the core PATH function *xcor* to calculate autocorrelations, both with default parameters.

### Lung Adenocarcinoma Metastasis

#### Dataset

Raw sequencing reads for RNA-sequencing and processed lineage data were downloaded from GEO (accession no. GSM4905334 and GSM4905335) for the M5K mouse. Aptly named because approximately 5K engineered A549 were initially implanted into the left lung of this immunodeficient mouse. To preprocess the raw RNA-sequencing reads, we used *scanpy* (Wolf, Angerer, and Theis 2018). Cells were initially filtered for total counts, number of genes with a positive count per cell, percent of total reads which are mitochondrial per cell, and the percent of counts in the top 20 genes. Per metric, we calculated an outlier cutoff based on 5 median absolute deviations, except for percent mitochondrial reads which has a cutoff of 3 median absolute deviations. Finally, we filtered genes for presence in a minimum of 20 cells. Altogether, this left just over 40K cells and 17,805 genes passing QC.

There were 81 trees publicly available for the M5K mouse. We pruned the trees to only include those cells with both lineage and gene expression using *ete3*.*pune()* function. We excluded trees with fewer than 32 cells after pruning for cells with both single-cell transcriptomics that pass QC metrics and lineage information which left 68 candidate trees. Tree sizes ranged from 33 cells to 9995 cells, with a mean size of 478 cells and a median of 90. For trees with sizes greater than 2000 (N=3), we down-sampled the trees randomly to 25% of their original size, maintaining relative tissue site proportions for left lung (LL), right lung (RL), mediastinum (M), and liver (Liv) cells. We included genes expressed in at least 20% of cells per tree in the test (average 3357 genes tested per tree [95% CI: 3195, 3529]). We used trees reconstructed from indel barcodes using the *Cassiopeia-Hybrid* pipeline (Jones et al. 2020; Huerta-Cepas, Serra, and Bork 2016).

#### Single-Cell Analysis

We subsetted the single-cell object to cells in tree LG13 and performed clustering using the Leiden community detection algorithm in Scanpy with resolution = 0.5 to generate 4 clusters. To identify differentially expressed genes, we used the *rank_genes_groups* function with method=’wilcoxon’ per Leiden cluster. We labelled any gene with an adjusted p.value <= 0.01 and abs(logFoldChange) > 1.5 in any cluster as being ‘differentially expressed’. We performed over-representation analysis as previously described using *C. elegans* using the MSigDB HALLMARK gene sets, which contain well defined biological states (Liberzon et al. 2015). We ran the *enrichR* function with a p-value cutoff of 0.05 and corrected for multiple testing using the Benjamini–Hochberg procedure.

## Data and Code Availability

The source code is implemented in *R* and is available at https://github.com/hrstuart/SCOUT. There is a simulation module to generate simulated data as described above and a model fit module. The Packer et al. *C. elegans* developmental dataset is available in the Gene Expression Omnibus (GEO) database with accession code <u>GSE126954</u> or through the *moscot* package (Klein et al. 2025). The lung cancer metastasis lineage dataset is available from GEO under accessions <u>GSM4905334</u> and <u>GSM4905335</u>.

## Acknowledgements

The authors would like to thank the members of the McKenna lab for their helpful discussions during the development of SCOUT. H.S. was supported by the Dartmouth Training Program in Quantitative Cancer Research (T32CA260262). This work was supported by N.I.H. award DP2GM149750 and A.M. is supported by the Pew Biomedical Scholars program.

---

**S1.**
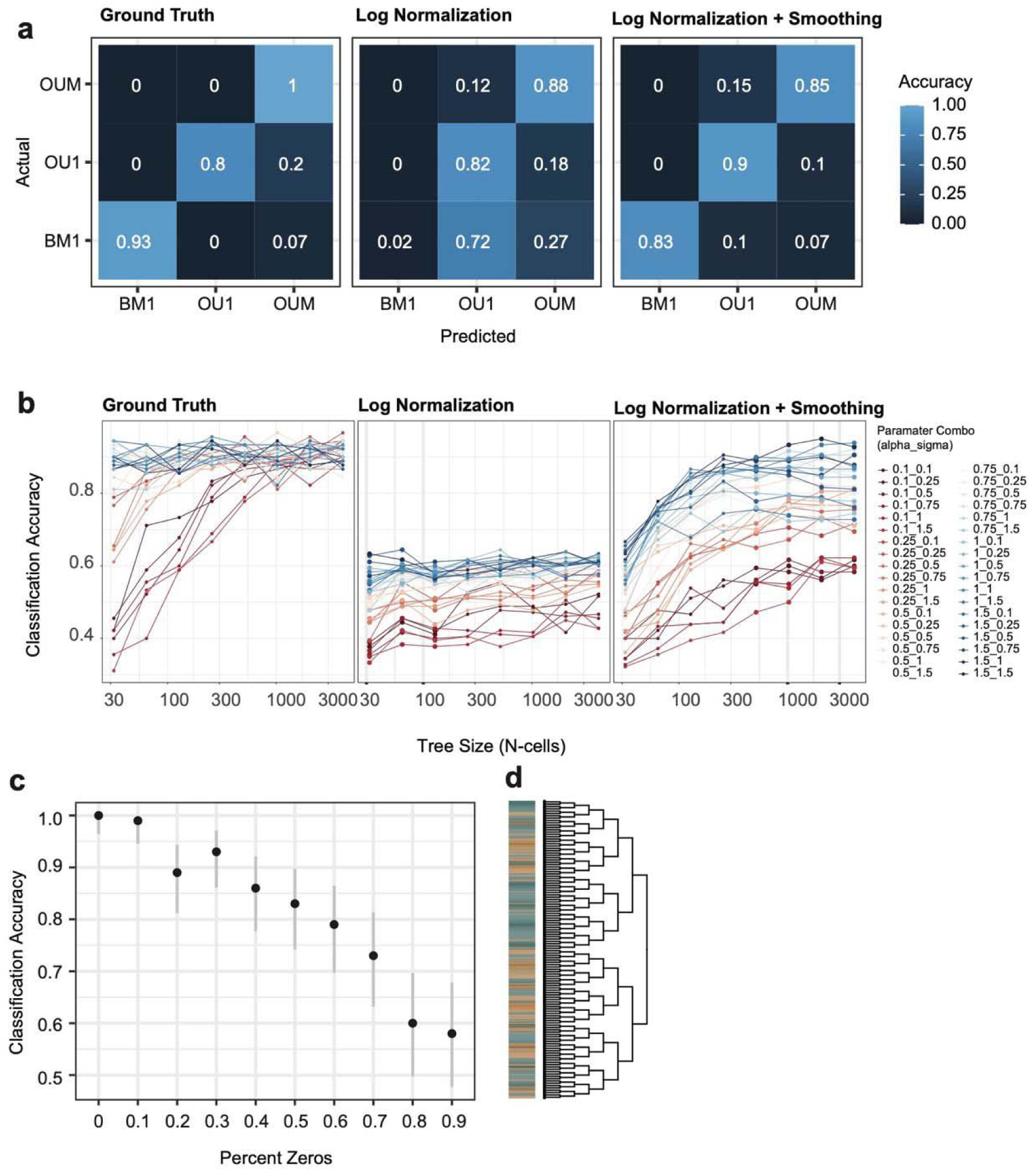
**Benchmarking SCOUT in simulated data**. (a) Confusion matrix illustrating the per class accuracy highlighting different strategies for preprocessing compared to a ground truth simulated example. The dataset was generated for n=512 cells, using alpha=0.75 and sigma=0.75. (b) Classification accuracy by tree size across parameters in a grid search, as seen in Figure 2D. Lines are colored by alpha magnitude (i.e. red lines have the smallest alpha value and dark blue lines have the largest alpha). (c,d) Classification results using a quasi-simulated example dataset in 293Ts such that ‘regimes’ were defined randomly assigned to one of two states (heatmap in d: state 1 in yellow and state 2 in green) and true gene expression values for highly variable genes were scaled to mimic an OU2 regime structure. Percent zeros are the number of zeros in the counts matrix, aiming to mimic drop-out.

**S2.**
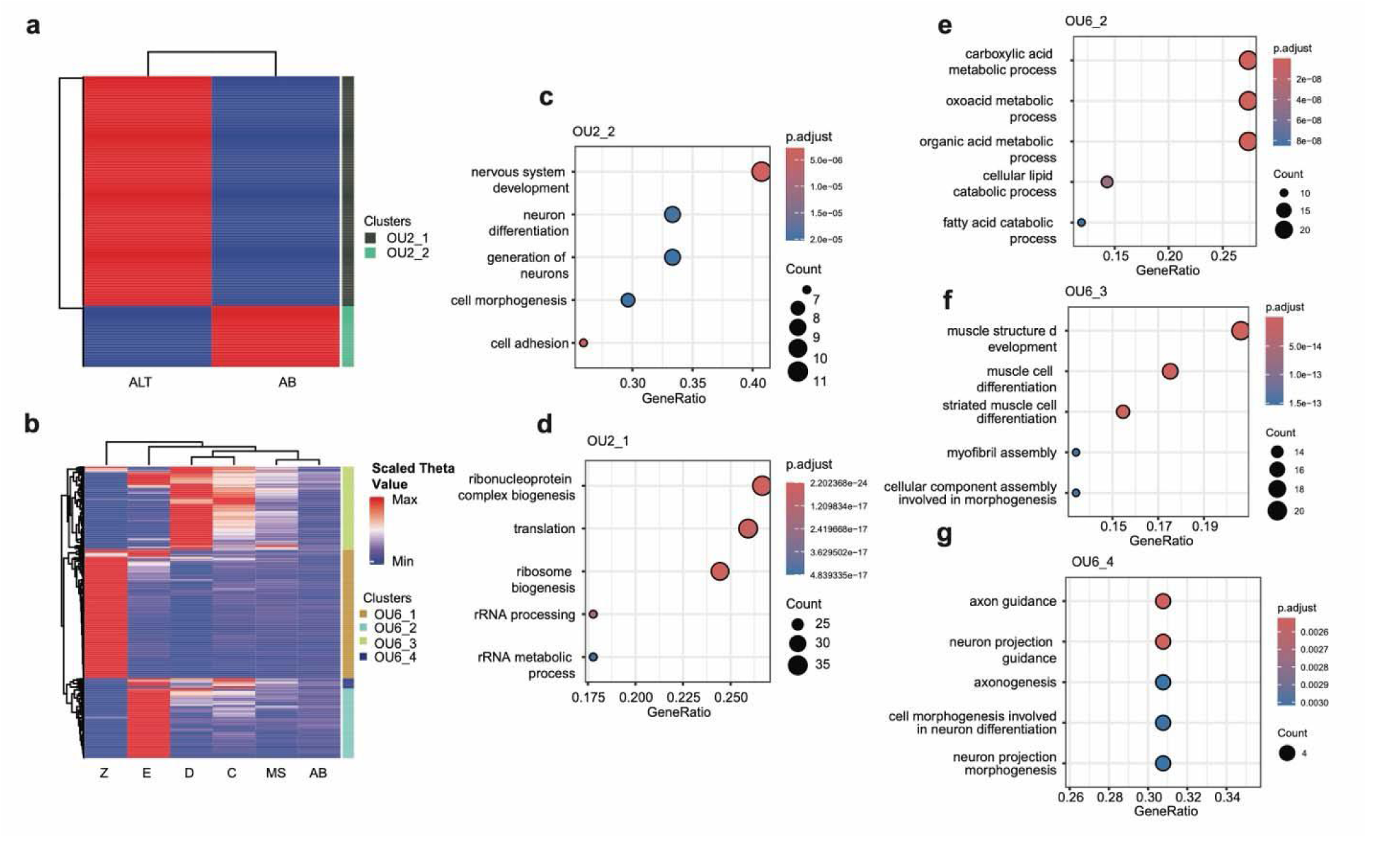
C. elegans gene ontology analysis. (a, b) Heatmap of theta optimal values for genes annotated as OU2 and OU6 (respectively). Clustering along the side is hierarchical clustering using Euclidean distance and cuttree to define clusters. Theta values were scaled using min/max scaling. (c-g) Gene ontology results using the GO term ‘aspect’ biological processes. (c and d) Top 5 GO terms for OU2 gene clusters and (e-g) top 5 GO terms for OU6 gene clusters. Three out of four OU6 gene clusters had valid GO terms.

**S3.**
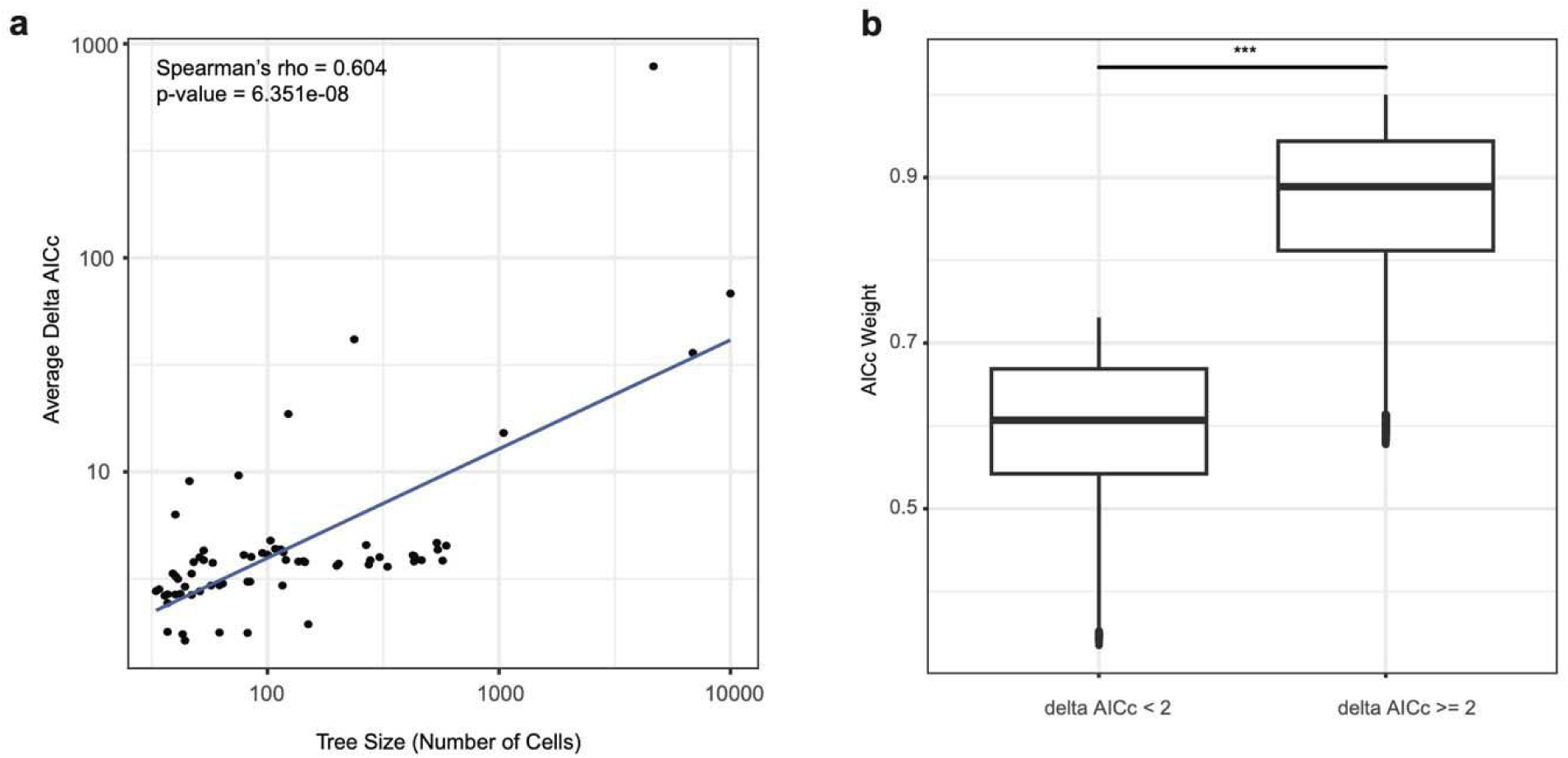
Performance of SCOUT in cancer metastasis datasets. (a) Spearman correlation between tree size and the average delta AICc to the next best model across all genes tested within a tree. (b) Comparing AICc weights between genes that did not pass delta AICc filter and genes that did.

**S4.**
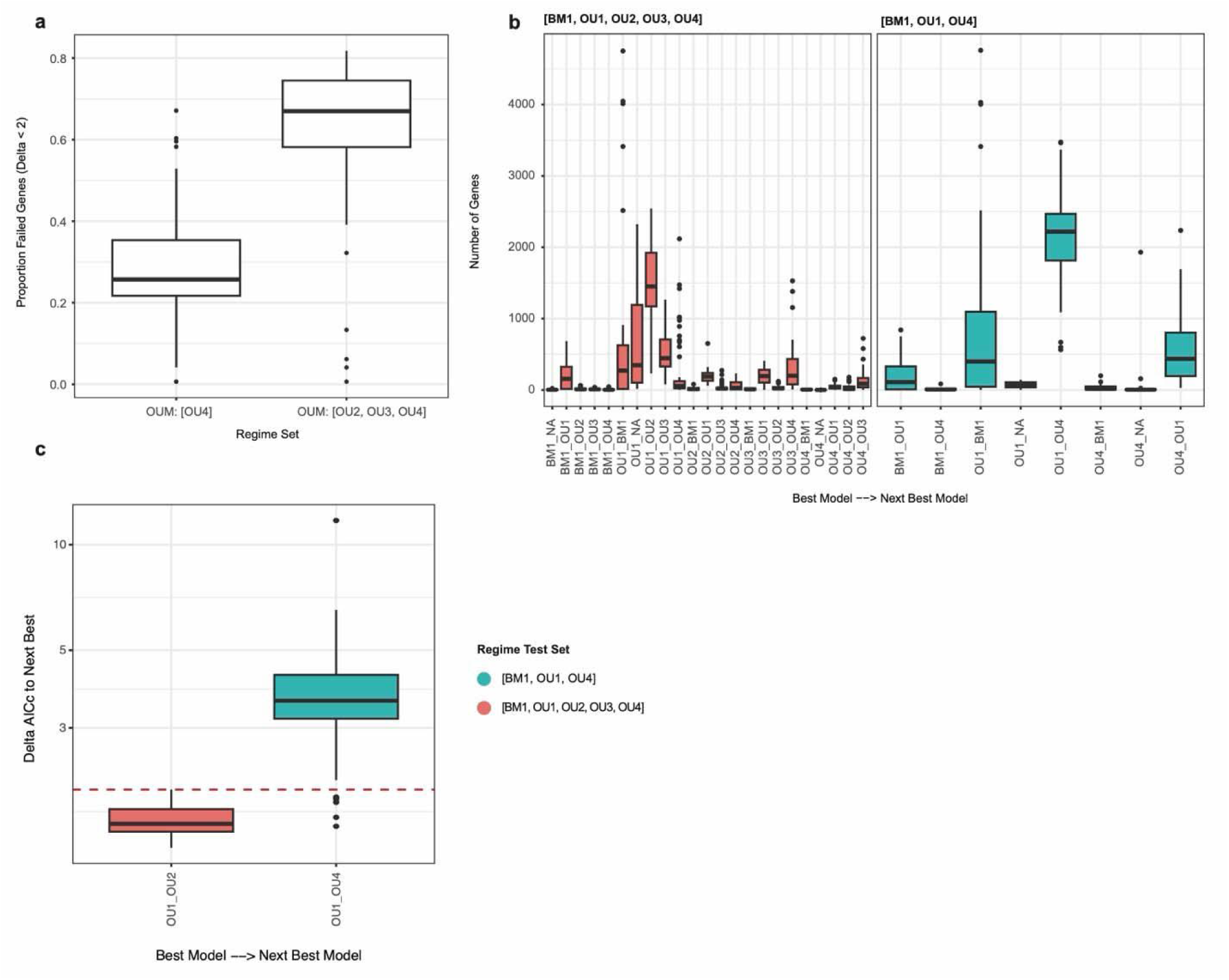
Selecting evolutionary hypotheses. (a) Across LUAD trees, two regime sets were tested: [BM1, OU1, OU4] and [BM1, OU1, OU2, OU3, and OU4]. Between the two sets, the boxplots report the distribution of the proportion of genes which fail to have a delta AICc of 2 to the next best model across all trees. (b) Comparing the frequencies of best-to-next-best models between the two regime sets. The boxplots reflect the distribution of the frequency across the tree set. (c) The distribution of delta AICc values for the best-to-next-best model pairs with the highest frequency across all trees, (OU1 v. OU2 for the full regime set, and OU1 v. OU4 for the reduced regime set). The horizontal red line indicates the delta threshold = 2.

**S5.**
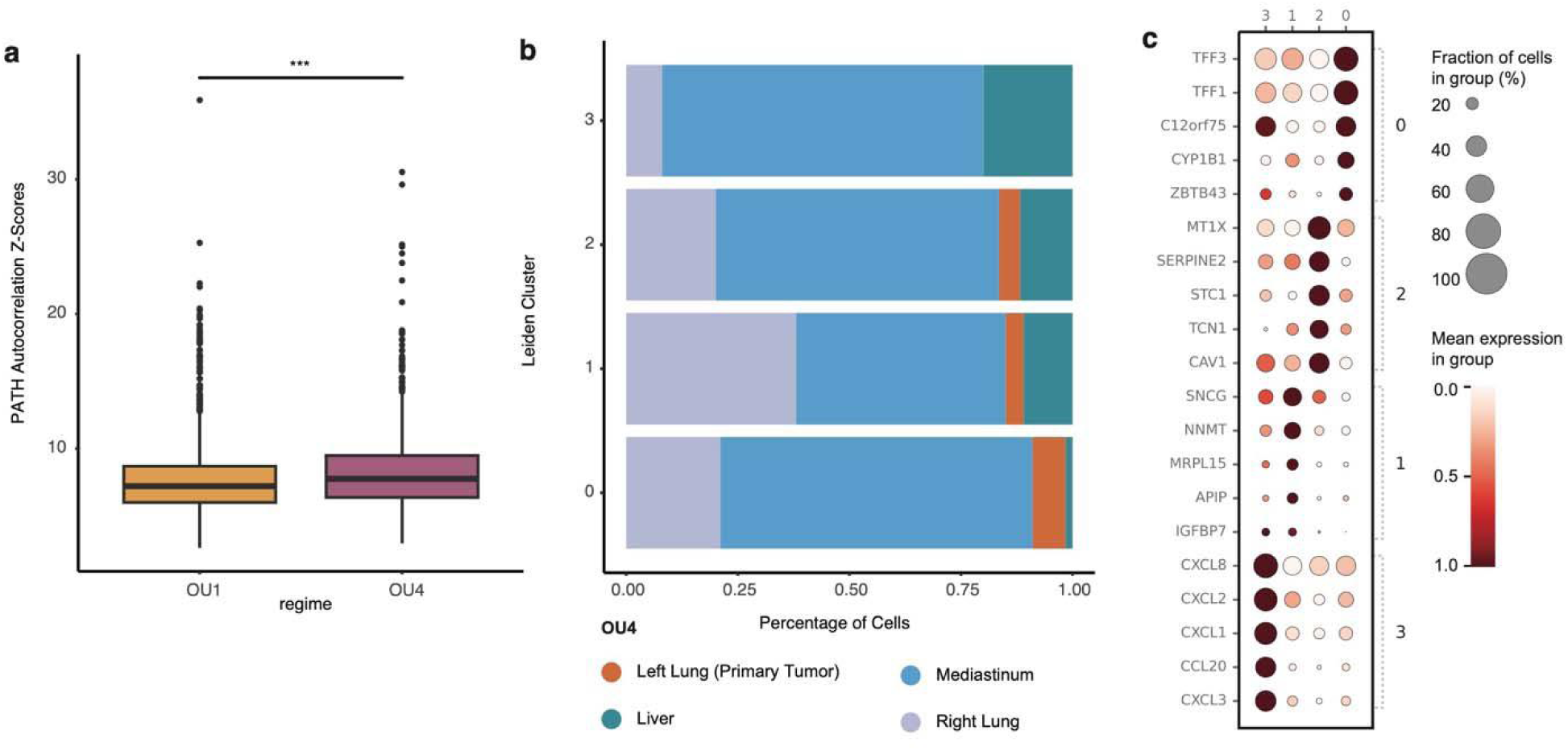
Comparing results to PATH and differential gene expression for LG13. (a) PATH autocorrelation Z-scores for OU1 and OU4 genes from LG13. (b) Proportion of cells from each metastatic site across the four Leiden clusters for LG13 single-cell UMAP (not shown). (c) Differential gene expression results for each leiden cluster. The top 5 genes in each cluster are included in the dotplot. Only genes with a log fold-change greater than 1.5 were included.

**S6.**
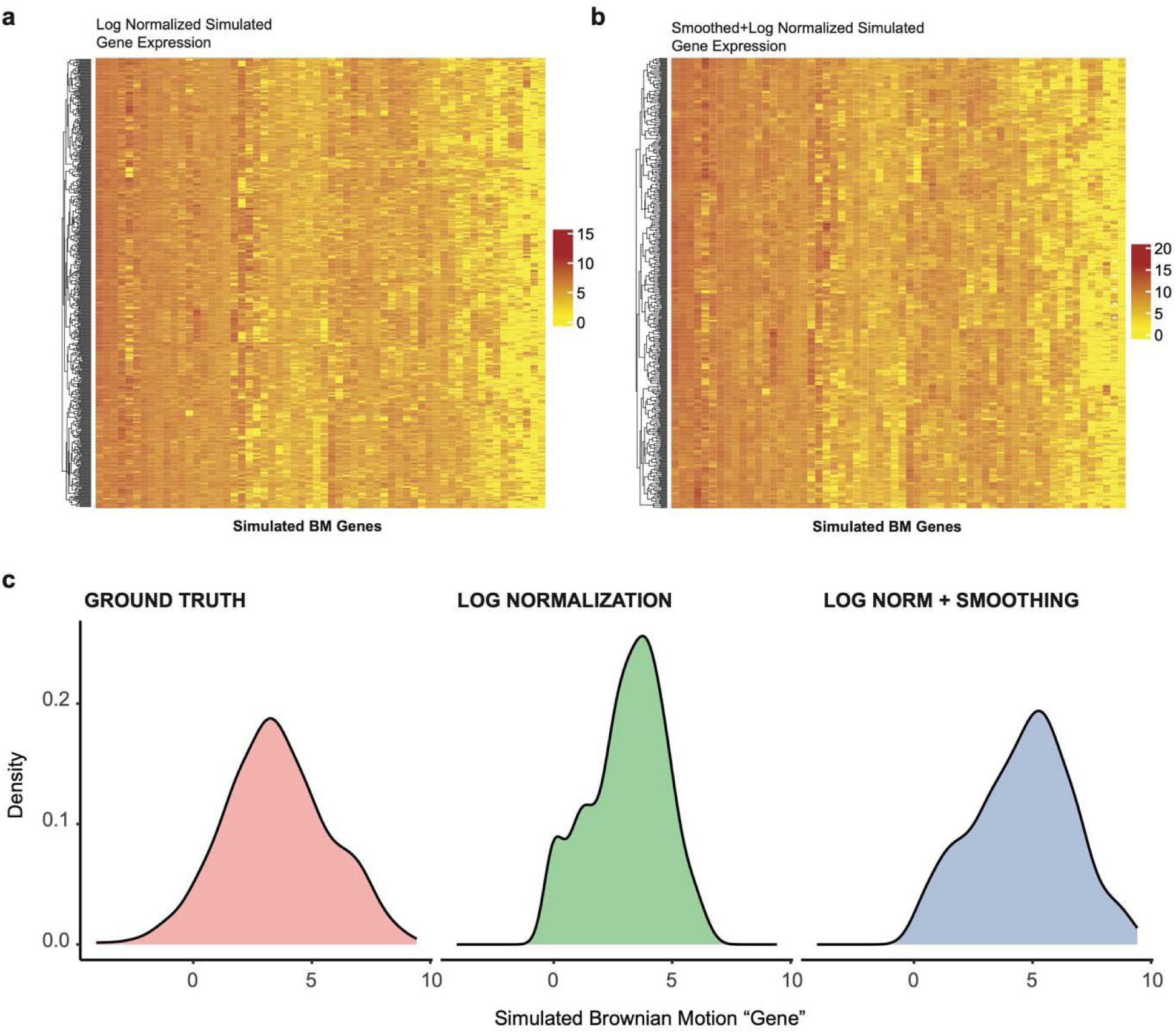
Example smoothing results from simulated data. Heatmaps illustrating simulated Brownian motion genes without (a) and with (b) smoothing. (c) Example distributions for a single gene comparing ground truth to log-normalized and log-normalized + smoothed data.

