## Supplemental Figures (S1-S6) for "SCOUT: Ornstein–Uhlenbeck modelling of gene expression evolution on single-cell lineage trees"

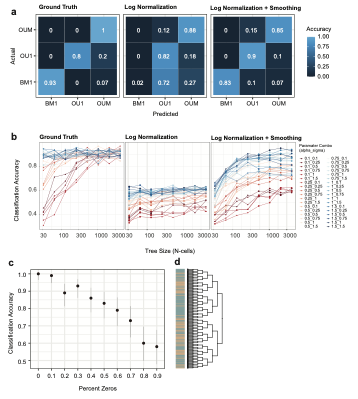

**S1. Benchmarking SCOUT in simulated data**. (a) Confusion matrix illustrating the per class accuracy highlighting different strategies for preprocessing compared to a ground truth simulated example. The dataset was generated for n=512 cells, using alpha=0.75 and sigma=0.75. (b) Classification accuracy by tree size across parameters in a grid search, as seen in Figure 2D. Lines are colored by alpha magnitude (i.e. red lines have the smallest alpha value and dark blue lines have the largest alpha). (c,d) Classification results using a quasi-simulated example dataset in 293Ts such that ‘regimes’ were defined randomly assigned to one of two states (heatmap in d: state 1 in yellow and state 2 in green) and true gene expression values for highly variable genes were scaled to mimic an OU2 regime structure. Percent zeros are the number of zeros in the counts matrix, aiming to mimic drop-out.

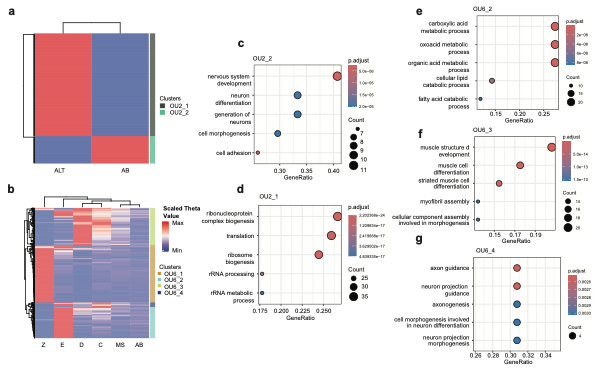

**S2. C. elegans gene ontology analysis.** (a, b) Heatmap of theta optimal values for genes annotated as OU2 and OU6 (respectively). Clustering along the side is hierarchical clustering using Euclidean distance and cuttree to define clusters. Theta values were scaled using min/max scaling. (c-g) Gene ontology results using the GO term ‘aspect’ biological processes. (c and d) Top 5 GO terms for OU2 gene clusters and (e-g) top 5 GO terms for OU6 gene clusters. Three out of four OU6 gene clusters had valid GO terms.

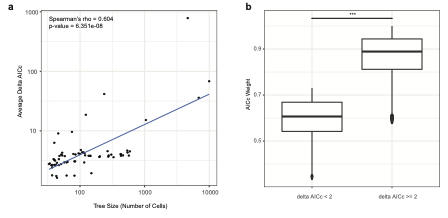

**S3. Performance of SCOUT in cancer metastasis datasets.** (a) Spearman correlation between tree size and the average delta AICc to the next best model across all genes tested within a tree. (b) Comparing AICc weights between genes that did not pass delta AICc filter and genes that did.

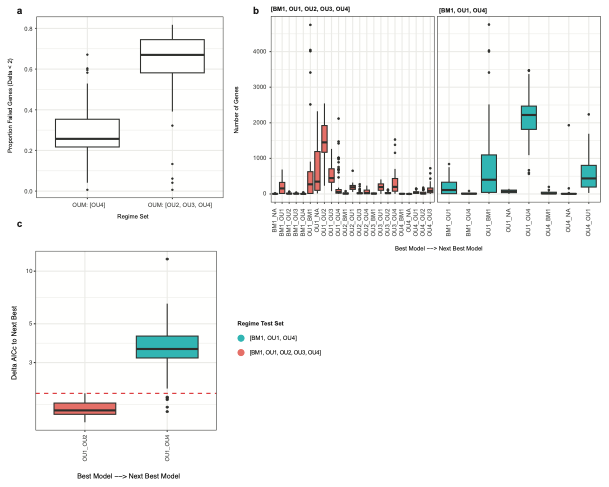

**S4.** **Selecting evolutionary hypotheses.** (a) Across LUAD trees, two regime sets were tested: [BM1, OU1, OU4] and [BM1, OU1, OU2, OU3, and OU4]. Between the two sets, the boxplots report the distribution of the proportion of genes which fail to have a delta AICc of 2 to the next best model across all trees. (b) Comparing the frequencies of best-to-next-best models between the two regime sets. The boxplots reflect the distribution of the frequency across the tree set. (c) The distribution of delta AICc values for the best-to-next-best model pairs with the highest frequency across all trees, (OU1 v. OU2 for the full regime set, and OU1 v. OU4 for the reduced regime set). The horizontal red line indicates the delta threshold = 2.

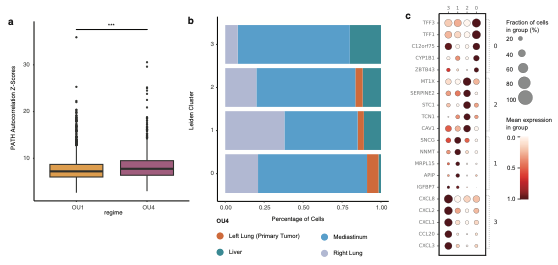

**S5. Comparing results to PATH and differential gene expression for LG13.** (a) PATH autocorrelation Z-scores for OU1 and OU4 genes from LG13. (b) Proportion of cells from each metastatic site across the four Leiden clusters for LG13 single-cell UMAP (not shown). (c) Differential gene expression results for each leiden cluster. The top 5 genes in each cluster are included in the dotplot. Only genes with a log fold-change greater than 1.5 were included.

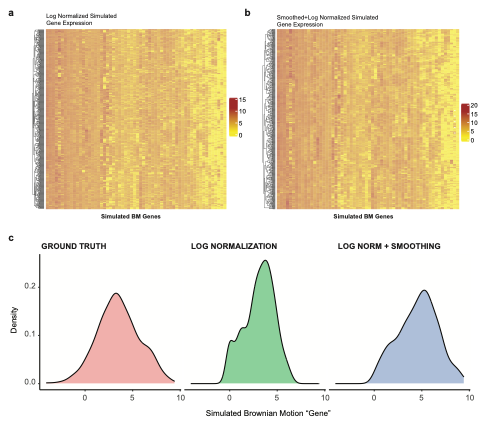

**S6. Example smoothing results from simulated data.** Heatmaps illustrating simulated Brownian motion genes without (a) and with (b) smoothing. (c) Example distributions for a single gene comparing ground truth to log-normalized and log-normalized + smoothed data.
